# Honest cues contribute to male choice for female guarding in a herbivorous spider mite

**DOI:** 10.1101/2021.09.07.459265

**Authors:** Steven F. Goossens, Frederik Mortier, Thomas Parmentier, Femke Batsleer, Thomas Van Leeuwen, Nicky Wybouw, Dries Bonte

## Abstract

Mate choice is a wide-spread phenomenon with important effects on ecological and evolutionary dynamics of successive generations. Increasing evidence shows that males can choose females if females vary in quality and these mating choices can strongly impact fitness. In the herbivorous spider mite *Tetranychus urticae* males engage in precopulatory mate guarding of quiescent females, and it is known that females vary in their time to sexual maturity and fecundity. However, our understanding of how males maximize their reproductive success and which female phenotypic traits are important cues for their mating decisions are still limited. In many arthropod species, female body size and pheromones are well known proxies for fecundity. These traits—and thus possibly male mating decisions—are however sensitive to environmental (dietary) stress. By allowing males to freely choose amongst many (synchronized) females in a controlled semi natural environment, we found that guarded females have a higher fecundity and are closer to sexual maturity than non-guarded females. Despite the fact that female body size was positively correlated with fecundity and significantly influenced by the environment, males did not discriminate on body size nor did we find evidence that they used other cues like cuticular pheromones or copying behavior (social cues). In conclusion we were able to show male mate preference for females that are closer to sexual maturity and have higher fecundity, but we were unable to identify the female traits that signal this information

## Introduction

In animals, mate choice can be defined as the specific behavior by which the quality of potential mates is assessed before copulation or fertilization (Bateson, 1983; Edward, 2014). The paradigm that females are a limiting factor in animal reproduction because of costly eggs (Bateman, 1948) is an erroneous simplification of complex eco-evolutionary processes (Edward & Chapman, 2011; Tang-Martínez, 2016). Males of many animal species provide nuptial gifts, invest in lengthy courtship or mate guarding, further reducing the number of future copulations (Kvarnemo & Simmons, 1998; Vahed, 1998; Byrne & Rice, 2006; Han & Jablonski, 2010; Papadopoulos et al., 2010; Oku, 2014). Such investment costs for males, favor male choosiness if female quality varies and this quality can be perceived by the males (Snowdon, 1997; Bonduriansky, 2001; Edward & Chapman, 2011; Hare & Simmons, 2019).

In many arthropods it is known that males prefer females with larger body size as it is associated with larger reproductive output (Honěk, 1993; Calvo & Molina, 2005; Drapela et al., 2013). Males use visual and/or tactile cues to evaluate female body size and adjust their mating behavior according to the estimated size (Gage, 1998; Byrne & Rice, 2006; Chenoweth et al., 2007). Chemical signals such as pheromones can signal mating status and sexual maturity and equally provide honest information about female fecundity (Gaskett, 2007; Steiger & Stökl, 2014). These chemical cues may be differentially perceived depending on environmental conditions or distances from the potential mate (Bel-Venner et al., 2008) . Besides gathering information about female quality directly, individuals may use social cues about potential mates as well. In this way, males can save time and energy by copying other individuals instead of thoroughly investigating the female quality themselves (Scauzillo & Ferkin, 2019). Such a choice copying is considered to be context-dependent (Witte, Kniel, & Kureck, 2015), leading to contrasting results in series of empirical studies with respect to the strength and reliability of female signals (Gaskett, 2007). This is because strong manipulative studies have their own constraints and introduce biasing effects, rendering insights on the relevance and significance of mate choice elusive. Many choice test, for instance, overestimated mate preference (Dougherty & Shuker, 2015).

We here aimed to avoid such constraints by testing mate selection in the two-spotted spider mite *Tetranychus urticae* under semi-natural conditions. *T. urticae* guard females that are about to molt to the adult stage (the quiescent female deutonymphs, or teliochrysalids). This guarding behavior allows males to secure the first mating opportunity and fertilize all the eggs of the guarded female (commonly referred as first sperm precedence; (Helle, 1967). Guarding is time-consuming and exposes spider mite males to predators, diseases and competitors (Alcock, 1994; Lima, 1998; Oku & Yano, 2008; Oku, 2009) and induces aggressive interactions among males (Potter, Wrensch, & Johnston, 1976) ,potentially result in wounding or even death (personal observations SG, NW; Potter, Wrensch, & Johnston, 1976). Therefore, males employing precopulatory mate guarding and fighting are anticipated to maximize their reproductive success by choosing the most fit females to guard. Males tend to select teliochrysalids that are close to emerge hereby minimizing the time invested per copulation (Parker, 1974; Everson & Addicott, 1982). Despite this intuitive appeal of the time minimization hypothesis, later studies generated inconsistent results in *T. urticae* (Saito, 2010). These non-consistent results may result from varying environmental conditions especially the population density and sex-ratio, as this directly affects the level of mate competition (Macke et al., 2012).

Consistent with studies that uncovered a limited visual capacity of *T. urticae* (Naegele, McEnroe, & Soans, 1966; W. D. McEnroe & Dronka, 1966; William D. McEnroe, 1969), chemical sensing of foliar structures and webbing have been implicated as important determinants in male searching behavior for female teliochrysalids (Cone et al., 1971; Penman & Cone, 1972). Some evidence on the importance of sex-pheromones has been provided in the 70ies of the previous century (Regev & Cone, 1975, 1976, 1980), but their effective function in signaling remains elusive and no volatile pheromones have been discovered that are known to contain information about the condition or quality of potential partners (Royalty, Phelan, & Hall, 1993; Oku, 2014; Oku et al., 2015). There is some evidence of mate choice copying behavior in spider mites, with males preferring guarded over solitary females, most likely because the former invest into a higher pheromone production (Oku & Shimoda, 2013). Paradoxically, solitary females had the highest fecundity, most likely because of being released from this costly pheromone production. This mating strategy therefore increases female fitness by attracting the most fit males, but not male fitness (Biernaskie, Grafen, & Perry, 2014). Given the fast resource exploitation dynamics of the species, dispersal rates (and therefore the potential to monopolize new resources) are an important fitness-related measure as well (Massol, Calcagno, & Massol, 2009; Bonte & Dahirel, 2017).

In this study, we observed mate guarding behavior of *T. urticae* under low and high population density in semi-natural conditions where the males can assess many different females before their mate choice. This allowed us to separate guarded and unguarded females under the different environmental conditions, and enables us to document putative alternative mating tactics of males. There is some evidence that body size is positively correlated with fecundity in this species (Li & Zhang, 2018). It has been shown that the CHC profiles of several insect species serve as indicators of female fertility as well (Thomas & Simmons, 2010; Bilen et al., 2013; Smith & Liebig, 2017), and that they may even be genetically correlated with fecundity (Berson & Simmons, 2019). In contrast to volatile pheromones, such CHC’s have not been investigated in spider mites. Lastly, we tested if males copy conspecific males by evaluating the spatial configuration of guarded females on the total leaf. We suspect that copying behavior and possible clustering of teliochrysalids could result in aggregations of guarded and unguarded females. We evaluated potential fitness maximization, and therefore the honesty of cues, by quantifying life history traits of all guarded and non-guarded females.

## Materials and Methods

### Mite husbandry

The *T. urticae* strain Lede was used in this study and is derived from the reference laboratory strain LS-VL that was originally collected from roses in Ghent, Belgium (Van Leeuwen, Stillatus, & Tirry, 2004). Mites of the stock population were reared on potted bean plants (*Phaseolus vulgaris* L. cv. ‘Prelude’) in a climate-controlled room at 26 ± 0.5 C, 60 % RH and 16/8 h (L/D) photoperiod. To create high (HD) and low (LD) densities, 40 (HD) and 10 (LD) adult females were randomly collected from the stock populations and transferred to leaf discs of 3.5 x 4.5 cm^2^, (freshly cut from 2-week-old plants) on wet cotton in an incubator (27 °C, 16:8 LD). Females were subsequently allowed to oviposit for 24 hours. On the 40 HD leaf discs there were an average of 15 eggs/cm^2^, whereas 40 LD leaf discs held on average 2 eggs/cm^2^. After 7 days left to develop in an incubator (37°C, 16:8 LD) most of the female mites were in the teleochrysalids stage (nymphal females in their final quiescent stage). On these leaf discs with females in the teleochrysalids stage, most male spider mites already reached adult hood and are present in a ratio of 1:3 (one male for every 3 females). Some males guard quiescent deutonymph females, the stage immediately before adult emergence and sexual maturation. The duration of the quiescent deutonymph stage is ca. 1 day at 27°C with a L16:D8 photoperiod (Kabiri, Saboori, & Allahyari, 2012). Immediately after emergence the guarding male quickly copulates with the adult female. The experiments were conducted in 2 batches, with a total of 35 HD,LD plates in the first set ad 35 HD, LD plates in the second experiment.

We let the males choose freely among many different teleochrysalids and subsequently measured their spatial clustering. Fitness related traits were measured by subsequently transferring individual guarded and unguarded teleochrysalids to bean leaf discs (2 x 2 cm^2^) that were connected by parafilm bridges on top of moist cotton and were lined with 2-mm-wide strips of moist tissue paper. For both the HD and LD treatments, 30 guarded and unguarded teliochrysalids were selected from random plates in both experiments, making a total of 240 individuals measured for body size, fecundity, survival and dispersal. After measuring the teliochrysalids body size the guarding male was introduced on the leaf disc to fertilize the emerging female, a random male was introduced on leaves of non-guarded teleochrysalids. In the second experiment no males were introduced on the leaf disc and thus all females are not mated. Time till emergence was tested for mites from the two batches of experiments, with respectively 38, 29 and 26 guarded females and 35,27 and 28 unguarded females for both experiments. Teleochrysalid cuticular chemical profiles were quantified from pools of 30 guarded and unguarded teleochrysalids from experiment 1.

### Spatial clustering of guarded females

We allowed all males from the age-synchronized populations to choose female teleochrysalids on the leaf discs and subsequently analyzed the spatial configuration of guarding males. If males were attracted by other males (through social cues), we expected a clustering of guarded teleochrysalids on the bean leaves.

To obtain the spatial configuration of guarding males, the seven day old leaves were photographed in a standard way of six leaves (3 HD and 3 LD). The positions of the guarded and unguarded teleochrysalids were marked using ImageJ (Schneider, Rasband, & Eliceiri, 2012) and were confirmed by manual inspection under a stereo microscope. After one hour, leaf discs were screened again, which confirmed that all females marked as guarded had remained guarded. The spatial positions were analyzed as point patterns with Spatstat (Baddeley, Rubak, & Turner, 2015). To analyze if guarded females were more clustered in the point pattern, the mark equality function is calculated for the six patterns. This function gives the probability that two points have the same mark (guarded or unguarded) when separated by a distance r. If guarded females are more clustered, this function would be higher at small distances than complete spatial randomness (horizontal line at value 1). Acceptance regions were drawn around the complete spatial random line using Monte Carlo simulation envelopes (n=500) to infer if the mark equality functions differ from complete spatial randomness.

### Fitness related traits in guarded and unguarded mites

#### Body size

we measured the teleochrysalid abdomen surface area (Troscianko, 2014) from high-resolution photos (Leica M50, 5 MP HD Microscope Camera Leica MC170 HD).

#### Fecundity

we tested whether male guarding was related to female reproductive performance by quanifying the number of offspring from a single females after 12 days (Wybouw et al., 2015).

Body size and fecundity were analysed by means of a linear mixed effect (lme) model in R using the package ‘lme4’. Mother density treatment, guarding status and their interaction were used a explanatory variables. We also included the iteration of the experiment as a variable intercept. Post-hoc analysis was performed on the estimated marginal means of the lme using the ‘emmeans’ package.

#### Survival

female mortality was monitored during a period of 20 days. We fitted a mixed-effect Cox model of how female mortality in time depended on whether the female was guarded or not. Females that died in the first 24 minutes were omitted because early deaths are attributed to the experimental manipulation. The mixed-effect Cox model enabled us to control for the possible auxiliary effects that were present during the two repetitions by including it as a variable intercept. We verified the proportional hazard assumption on each experiment separately (Supplementary statistics ,2).

#### Time till emergence

We tested whether males could perceive time to emergence when deciding which female to guard. Teleochrysalids that were guarded or unguarded for at least one hour (see marking for the spatial analysis) were individually transferred to bean leaf square (4 x 4 cm^2^). The leaf was divided in two half’s and guarded females were placed on one side and the unguarded on the other. A series of time laps pictures was taken (Leica M50, 5 MP HD Microscope Camera Leica MC170 HD) every 10s for 24h. The pictures of the time laps were analyzed to determine the molting time of each guarded and unguarded female.

We fitted a mixed-effect Cox model of how female emergence in time depended on whether the female was guarded or not using the ‘survival’ and ‘coxme’ packages (Therneau et al., 2021). The mixed-effect Cox model enabled us to control for the possible auxiliary effects that were present during the three repetitions by including it as a variable intercept. Data from the first experiment violated the proportional hazard assumption and were excluded from the formal analyses (but provided in supplement statistics,1a,1b).

#### Dispersal

We quantified dispersal by the movement of mites across parafilm bridges to adjacent leaf discs (Dahirel et al., 2019) for adult females in the first batch of experiments. Spider mite movements were monitored 4 times/day. We constructed a Kaplan-Meier model of how dispersal in time depended on whether females were previously guarded or not and on their density during testing. The survival and dispersal analyses were conducted with the ‘survival’ and ‘coxme’ packages (Therneau et al., 2021).

### Chemical profile of guarded and non-guarded females

Here, we wanted to test whether the teleochrysalid cuticular chemical profile affected male guarding behavior. We transferred pools of 30 guarded and unguarded teleochrysalids with hexane-cleaned brushes to separate 2-mL glass vials in five replicates (Sigma-Aldrich). In addition, we also transferred pools of 30 adult males to separate 2-mL vials in two replicates. The teleochrysalids and adult males were frozen and kept at −21 °C until solvent extraction and GCMS analysis. We extracted the cuticular compounds for 10 min in 2-mL vials capped with a PTFE septum (Sigma-Aldrich) in 20 μL of hexane (HPLC grade, Sigma-Aldrich). The hexane extract was transferred to another vial. Samples were left to evaporate at room temperature in a laminar fume hood and stored at −21 °C prior to analysis. Samples were diluted again in 20-μL hexane. We injected 2 μL of each hexane extract into a Thermo GC (Trace 1300 series) coupled with a MS (ISQ series, −70 eV, electron impact ionization) and equipped with a Restek RXi-5sil MS column (20 m × 0.18 mm × 0.18 μm). Extraction, identification and quantification of the CHCs was conducted as described in (Parmentier et al., 2018).

## Results

### Spatial clustering of guarded females

Mark equality plots of the three HD and three LD point patterns showed that guarded (or unguarded) females are not more clustered together than expected under complete spatial randomness (Supplementary fig. S1).

### Fitness related traits in guarded and unguarded mites

Female body size did not differ significantly between the guarding status (247±14.6μm for unguarded individuals, 243±14.6μm for guarded individuals, difference: 3.51μm, p = 0.1688, supplementary fig. S2). As expected from the more stressful high density treatment, female mites exposed to a high density were significantly smaller (t(density) = 4.692, p=<0.0001) than those mites kept in low density (237±14.5μm for HD individuals versus 253±14.6μm for LD individuals, difference: −16.1μm, p<0.0001, Supplementary fig.S2). The large difference in body size between HD and LD mites and the variation within the treatments shows that a variation in body size can be found, but in our experiment males did not prefer larger females.

Body size was positively correlated to fecundity (r=0.30; p= 0.00018, Supplementary fig. S3). The total number of offspring (fig. 1, panel a) differed between density treatments and between the guarding status of the female. Females were estimated to produce 63.2 ±3.78 offspring on high density leaves versus 73.1 ±3.82 offspring on low density leaves (difference: −9.98, p<0.0001, fig. 1, panel c). Furthermore, guarded females produced 71.6 ± 3.78 offspring, unguarded females 64.7± 3.79 offspring after 12 days (difference: 6.83, p = 0.0014, Fig. 1,panel b). We did not observe a significant interaction effect between mother density and guarding.

**Figure 1:**
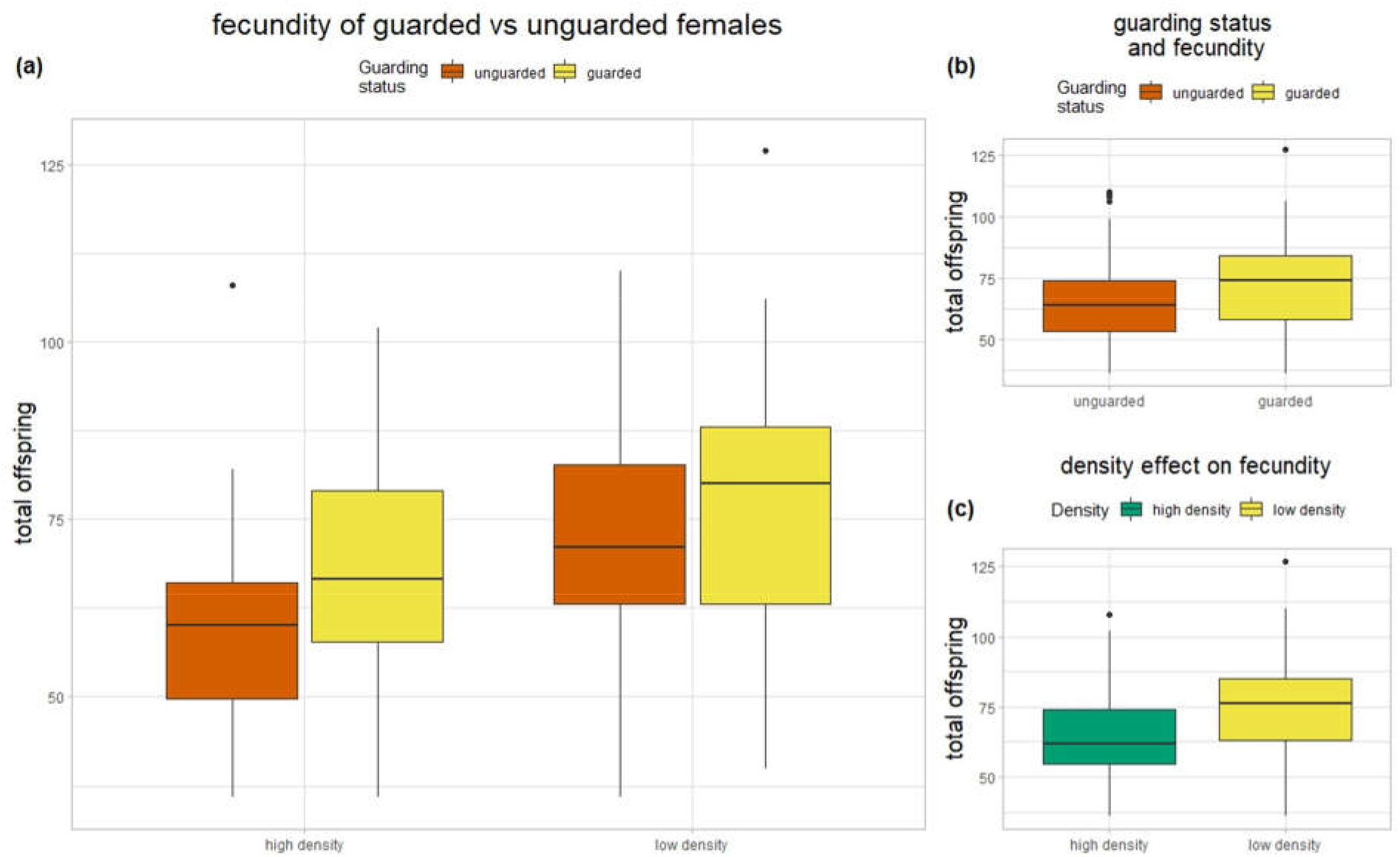
Fecundity of guarded females and unguarded females in high and low density plots. (a) fecundity (total offspring) of the guarded females is significantly higher in high (food stress) and low density conditions. (b) Total difference in fecundity of unguarded vs guarded females. (c) Total difference in fecundity between high density (food stress) and low density conditions.

Over both analyzed experiments, guarded teliochrysalids molted to adult females with a probability that was 1.82 higher than that of unguarded females (Cox mixed-effects model, exp(coeff) = 1.82, p=0.0023; Fig 2). Before 900 min (15h) we see at each time point that a higher proportion of guarded females emerged but after 900 min this levels out. Males seem therefore able to detect females that will emerge from their final ecdysis within 900 min.

**Figure 2:**
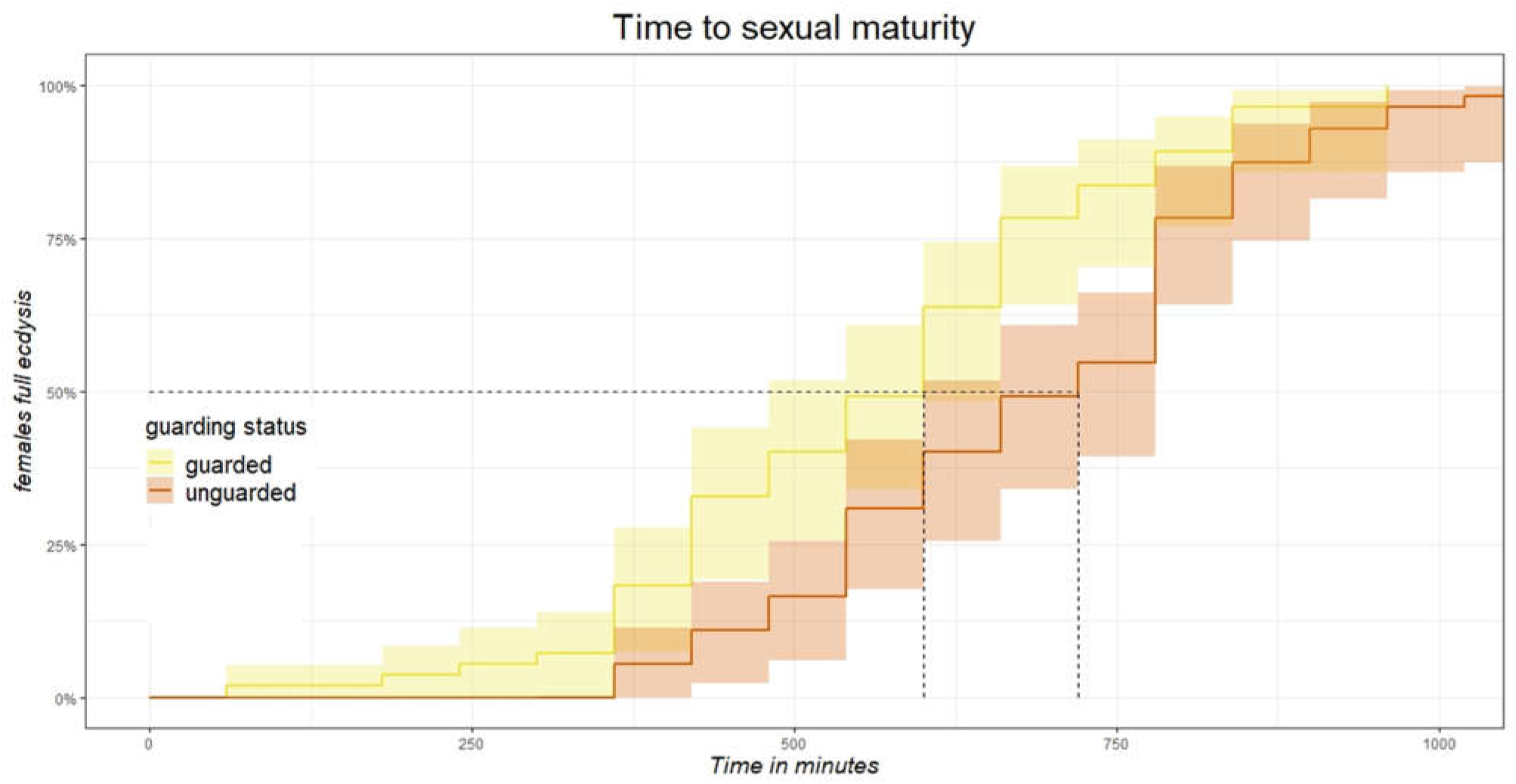
Time to event (sexual maturity) plot of guarded and unguarded females. Guarded females emerge faster from there resting state than unguarded females with the largest differences at the early time points (faster emergence) and less difference past 800 minutes (33h).

The Kaplan–Meier estimator shows that there were no differences in survival (p=0.48), or dispersal (p=0.75) between guarded and non-guareded females (see supplement statistics,2a,2b,3).

### Chemical profile of guarded and non-guarded females

The cuticular hydrocarbons of the spider mites constituted a homologous series of saturated straight chain alkanes from n-C22 to n-C33 (supplementary Figures,S4. There were quantitative differences in the CHC composition between males and females, males formed a separate group in an nMDS plot (Permanova—adonis, 999 permutations, Pseudo-F = 7.77, R^2^ = 0.6334, P = 0.008). but guarded females could not be separated graphically in an nMDS plot (supplementary Figures,S4). Consequently, no effect of treatment group on female CHC composition could be demonstrated (Permanova—ADONIS, 999 permutations, Pseudo-F = 1.04, R^2^ = 0.112, P = 0.439).

## Discussion

In this study, integrated experiments show that *T. urticae* males preferred to guard females that differed in sexual maturation and fecundity from unguarded females. Contrary to our hypotheses we did not find any evidence that males use cuticular hydrocarbons, body size or social cues in sexual selection.

As in other species (e.g., amphipods; Dick & Elwood, 1990), spider mite males likely rely on cues that provide information on both fecundity and time till maturity to enhance their mating success. Alternatively, mate guarding could shorten the female quiescence, so advance maturation in females rendering the presence of males rather the cause than the consequence of life history changes. Indeed, as for amphipods (Ward, 1984), spider mites females have been shown to delay molt in absence of mating partners (Oku, 2016).

We found that guarded females were more likely to be closer to molting than unguarded females with indications that small differences in molting in the last 15h to molting are detected. Conversely, Oku & Saito (2014) also found that males preferred females close to molting but could only discriminate if the time to molting was more than 22h apart. This can be explained by the fact that males in our experiment could assess all possible options, and reject females closer to maturation in favor of females that have the energy and resources to lay more eggs but take a bit longer to molt. Also some females close to maturation could already be guarded by other males, so males have to waste time and energy for male combat to take over the most optimal females or search for the next best female that is not guarded. By using both the time they have to invest in guarding and female fecundity, males can maximize their reproductive gains per unit time of investment. It is known that males usually do not spend much time in inter-male conflicts (<5s), with the approaching male being attacked and driven off by a guarding male in most cases (Royalty, Phelan, & Hall, 1993). In our experiment we also found that once males become arrested near a female, they tend to commit to guard that female until they mate with the guarded female. Alternatively, males can also vary in their preference and ability with no single female “type” favored by all males. It would therefore be interesting to study male traits in association with their choice and competition for specific female (traits). Moreover, we argue that male mate choice may be a threshold response where male investment is conditional to the available energy budget, density of females, other males, the time until the female reaches sexual maturity and total number of offspring per female.

The positive correlation between body size of the teleochrysalids and fecundity was rather modest (0.3) and males in our experiment did not prefer or could not distinguish the larger females. We did find that in the lower stress condition (low density), females were significantly larger than females raised in a stressful environment (high density). However, male choice for body size was not influenced by the environment. Similarly, females in the low stress conditions had significantly more offspring than the females in the stressful environment. Although males generally prefer larger females in many insects and spiders (e.g., Saeki, Kruse, & Switzer, 2005), females may also “chose” to allocate resources to reproduction instead of other traits such as growth and have therefore body size decoupled from fecundity. In these, large body size is not selected for (e.g., Wolz et al., 2020) or even counter-selected (e.g., Klingenberg & Spence, 1997).

A positive relation between fecundity and cuticular hydrocarbons (CHC’s) is found in many insects and quantitative and qualitative differences in CHC’s are important in for partner choice in spiders (Gaskett, 2007; Chung & Carroll, 2015; Berson & Simmons, 2019). In spider mites, volatile pheromones have long been investigated as an attractant for male spider mites, with sometimes contradicting results (Oku, 2014). Even though it is possible that males are attracted by volatile pheromones, it seems unlikely that these volatiles fond in spider mites so far provide information about the quality of the female (Saito, 2010; Oku et al., 2015). As in spiders we suspected that airborne sex pheromones typically attract males, but rarely elicit courtship (Gaskett, 2007). In contrast CHC’s not only provide males with information about female fecundity but in many species their composition also changes with sexual maturation (Ala-Honkola et al., 2020). Although previous studies used proxies to investigate the attraction of male spider mites to CHC’s and distinguish these from the attraction to volatiles (Rodrigues et al., 2017), no studies on their role as a possible cue for female quality exist. We found that cuticular CHC’s composition of *T. urticae* consists of a homologous series of saturated straight chain alkanes from c22 to c33. A similar result of CHC’s arranged in a series of straight chain alkanes was found in oribatid mites (Acari: Oribatida) (Raspotnig et al., 2008). We could not find any differences in quantity nor quality of the CHC’s between guarded and unguarded females. As a consequence of small sizes of the spider mites, and thus, minute amounts of hydrocarbons, a high number of individuals was necessary to prepare crude extracts with detectable alkane-profiles. This could also mean that a more subtle signal of female quality might not be noticed by our methods as we cannot evaluate every female separately. Alternatively, pheromones signaling female quality and sexual maturation could also be present in the webbing of the resting females as is known form spiders (Baruffaldi et al., 2010; C. Scott et al., 2018). Although it is suggested that spider mites can use silk as a movement cue, nothing is known about what information is present in the silk (Penman & Cone, 1974). For instance during web production female spider can deposit large amounts (up to 5μg/web) of pheromones in their silk, providing possible mates with cues about their condition (Schulz & Toft, 1993; Gaskett, 2007).

Finally, we did not find any evidence that males use social cues in guarding behavior. Although there were multiple teliochrysalids close to each other, mostly only one or two of these were guarded. The use of social information in mating is not uncommon in spiders and insects (Mery et al., 2009; Fowler-Finn et al., 2015; C. E. Scott, McCann, & Andrade, 2019). Relying on social information, mate-choice copying, and thus aggregating close to deceptive females, is costly as it increases conflict and direct mortality (Potter, Wrensch, & Johnston, 1976). Hence, male mate-choice copying behavior is only expected to occur when searching (time) costs are high (Real 1990). In spider mites, searching times are likely not very high as mating partners are quite densely distributed. This contrasts strongly with systems where such behavior was observed, e.g. in spiders where mate searching can have high mortality rates, favoring efficient detection of and movement towards females (C. E. Scott, McCann, & Andrade, 2019). Our results indicate that males avoid to guard females that are attended by a number of males. We cannot rule out that there was conflict between males before they settled in their final choice, but in all cases there were still more females unguarded than guarded, leaving ample variation. We neither found any adjustment of these mate selection strategies over the two density treatments, despite changes in phenotypes.

Although pervious research found that spiders mites changed their guarding behavior in response to density we did not find any indication that males changed their guarding behavior (Oku, 2009). We did find that the body size and fecundity differed for the females between our density treatments indicating an effect of the stressful conditions. Nevertheless it is possible that our ‘high density’ was not high enough for the males to alter their guarding behavior.

In conclusion, we here demonstrated non-random mate selection of spider mites in semi-natural settings. Our work complements earlier work using choice experiments and shows that even in conditions where multiple cues are present, that sexual selection is based on honest cues, i.e., that males select females directly providing largest fitness benefits by selecting females that minimize time costs (guarding time) as well as maximizing reproduction. This signaling appears to be based on other cues than cuticular hydrocarbons, body size or social information.

## Supporting information

supplementary statsistics to Goossens et al. 2021

Supplementary figures to Goossens et al. 2021

## Funding and Acknowledgements

Steven F. Goossens is funded by Research Foundation - Flanders (FWO) (FWOproject G018017N). Frederik Mortier and Femke Batsleer are funded by FWO. NW was supported by a BOF post-doctoral fellowship (Ghent University, 01P03420)

We thank the bachelor students Ditte De Waele and Lennard Dekeukeleire and lab assistant Ruben Claus for their contribution in gathering the data for this research.

## Author contributions

The study was initially conceptualized by S.G. and developed with input from D.B. and T.P. .S.G developed the experiments and collected data. T.P. extracted the cuticular hydrocarbons (CHC’s) and conducted gas chromatography and subsequent identification and statistical analysis of the CHC’s. S.G and F.M. performed the statistical analysis of life history traits and body size. FB performed the spatial analysis. S.G. and N.W. drafted the manuscript. All authors proofread the manuscript and made valuable contributions.

