## supplementary statsistics to Goossens et al. 2021 for "Honest cues contribute to male choice for female guarding in a herbivorous spider mite"

### Supplementary Statistical analysis

#### 1: Female emergence analysis

##### a. Test assumption of proportional hazard

We tested the proportional hazard assumption in R (3.6.3) using the cox.zph function of the ‘survival’ package that makes use of the conventionally-used Shoenfeld residuals. We tested this assumption for every iteration of the experiment separately.

|  | Chisq | Df | P |
| --- | --- | --- | --- |
| Experiment 1 | | | |
| Guard | 5.01 | 1 | 0.025 |
| GLOBAL | 5.01 | 1 | 0.025 |
| Experiment 2 | | | |
| Guard | 0.287 | 1 | 0.59 |
| GLOBAL | 0.287 | 1 | 0.59 |
| Experiment 3 | | | |
| Guard | 0.0252 | 1 | 0.87 |
| GLOBAL | 0.0252 | 1 | 0.87 |

Table S1: Proportional hazard assumption test for female emergence in all three experiments

In the first experiment, the variable that indicated whether the female was guarded (‘guard’) deviated significantly from the proportional hazard assumption (p = 0.025, table S1). In the other two experiments the assumption held. Therefore, we analyzed only the last two experiments in the mixed-effect Cox model.

##### b. Analysis of experiment 1

Here, we analyze experiment 1 using a Kaplan-Meier model of how female emergence in time depended on whether the female was guarded or not. This model does not make the assumption of proportional hazards.

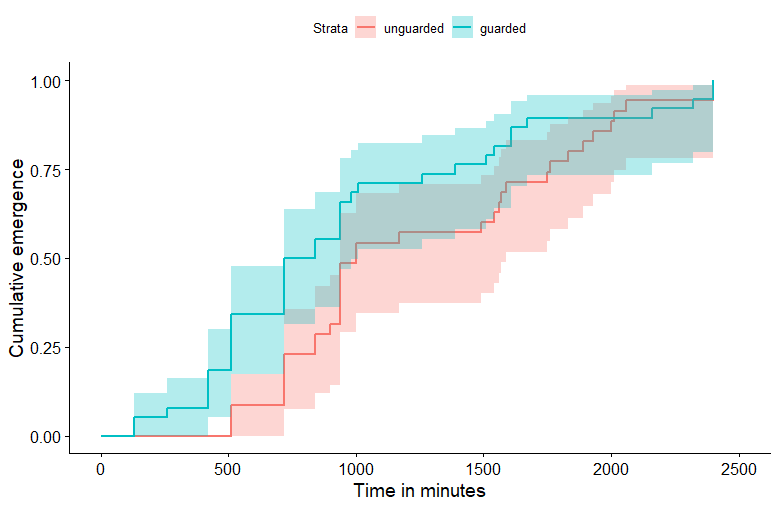

Figure S1: Survival plot of cumulative female emergence in experiment 1 using Kaplan-Meier with guarded (blue) and unguarded (red) females. Shaded areas indicate 95% confidence interval.

The survival curves show that guarded females tend to emerge more likely than unguarded females during most of the experiment with quite some overlap in confidence intervals (fig. S1). This is reflected in the lack of, but close to, significance in the survival difference test (table S2).

#### 2: Female mortality analysis

##### a. Test assumption of proportional hazard

A Cox proportional hazard model, and by extension a mixed-effect Cox model, assumes that the tested explaining variables have a proportional effect on the ‘hazard’, i.e. the probability of an event such as adult emergence happening, and that that proportion is constant in time. We tested this in R (3.6.3) using the cox.zph function of the ‘survival’ package that makes use of the conventionally-used Shoenfeld residuals. We tested this assumption for every iteration of the experiment separately.

|  | Chisq | Df | P |
| --- | --- | --- | --- |
| Experiment 1 | | | |
| Guard | 1.85 | 1 | 0.17 |
| GLOBAL | 1.85 | 1 | 0.17 |
| Experiment 2 | | | |
| Guard | 0.178 | 1 | 0.67 |
| GLOBAL | 0.178 | 1 | 0.67 |

Table S2: Proportional hazard assumption test for female mortality for experiments 1 and 2.

In both experiments the assumption of proportional hazards held (table S2).

##### b. Results

Over both the life-history experiments, guarded and unguarded females showed no significant difference in their mortality risk in time (Cox mixed effect model, exp(coeff) = 1.21, p = 0.18, fig. S2)

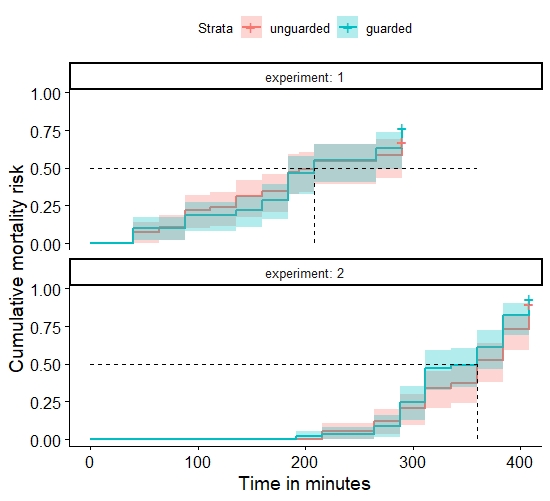

Figure S2: Predicted cumulative mortality function for a Cox proportional hazard function unguarded (red) and guarded (blue) females. Shaded areas indicate 95% confidence interval.

##### 3: Dispersal rate analysis

The Kaplan–Meier estimator shows that there were no differences in dispersal probability in time between guarded and unguarded females in high and low density settings (Chisq = 1.3, df = 3, p=0.7, fig. S3).

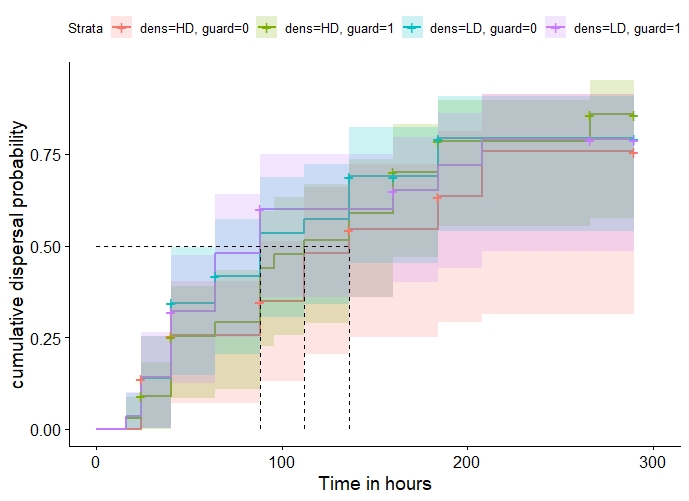

*Figure S3: Survival plot of cumulative dispersal probability using Kaplan-Meier with high density, unguarded (red); high density, guarded (green); low density, unguarded (cyan) and low density, guarded (purple) females. Shaded areas indicate 95% confidence interval.*
