## Supplementary figures to Goossens et al. 2021 for "Honest cues contribute to male choice for female guarding in a herbivorous spider mite"

**S1**

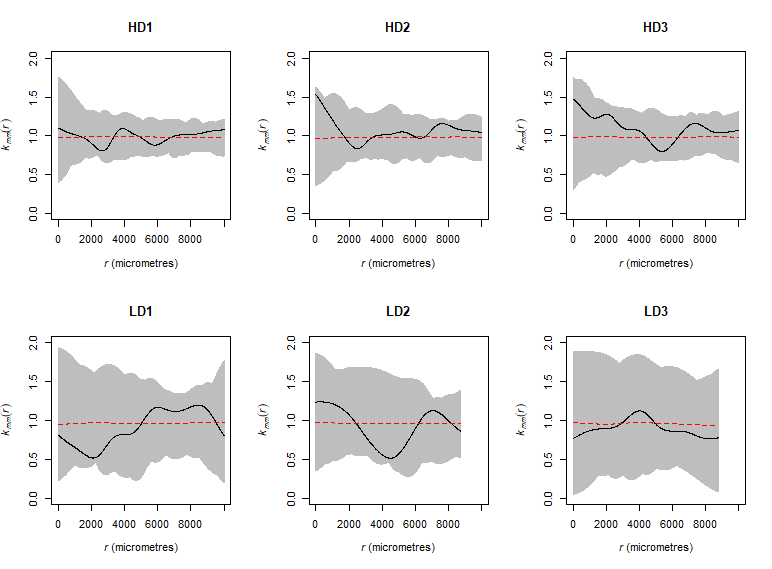

**Supplemental figure 1:** Mark equality plots of the three HD (upper panel) and three LD point patterns (lower panel) showed that guarded (or unguarded) females are not more clustered together than expected under complete spatial randomness.

**S2**

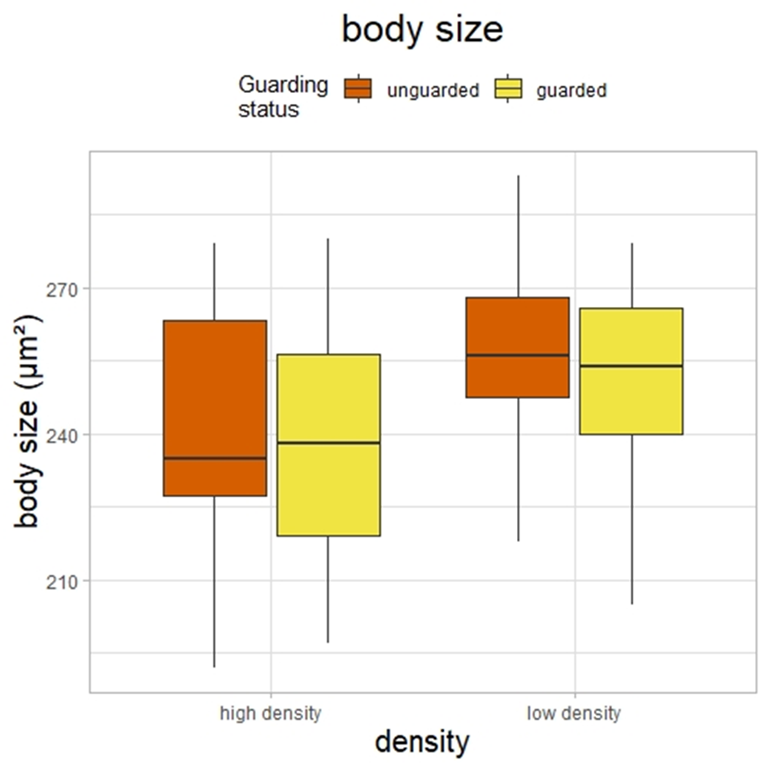

**Supplemental figure 2:** Body size of guarded and unguarded females in food stress (high density) and low stress conditions (low density). There is no difference in body size between the guarded and unguarded females. There is a significant difference in female body size between the high and low density conditions.

**S3**

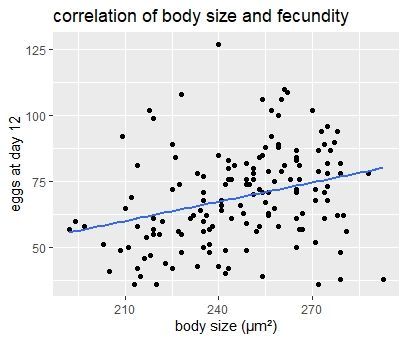

**Supplemental Figure 3**: Correlation of the total offspring at day12 (y-axis) and body size of the female teleochrysalids (x-axis).

**S**
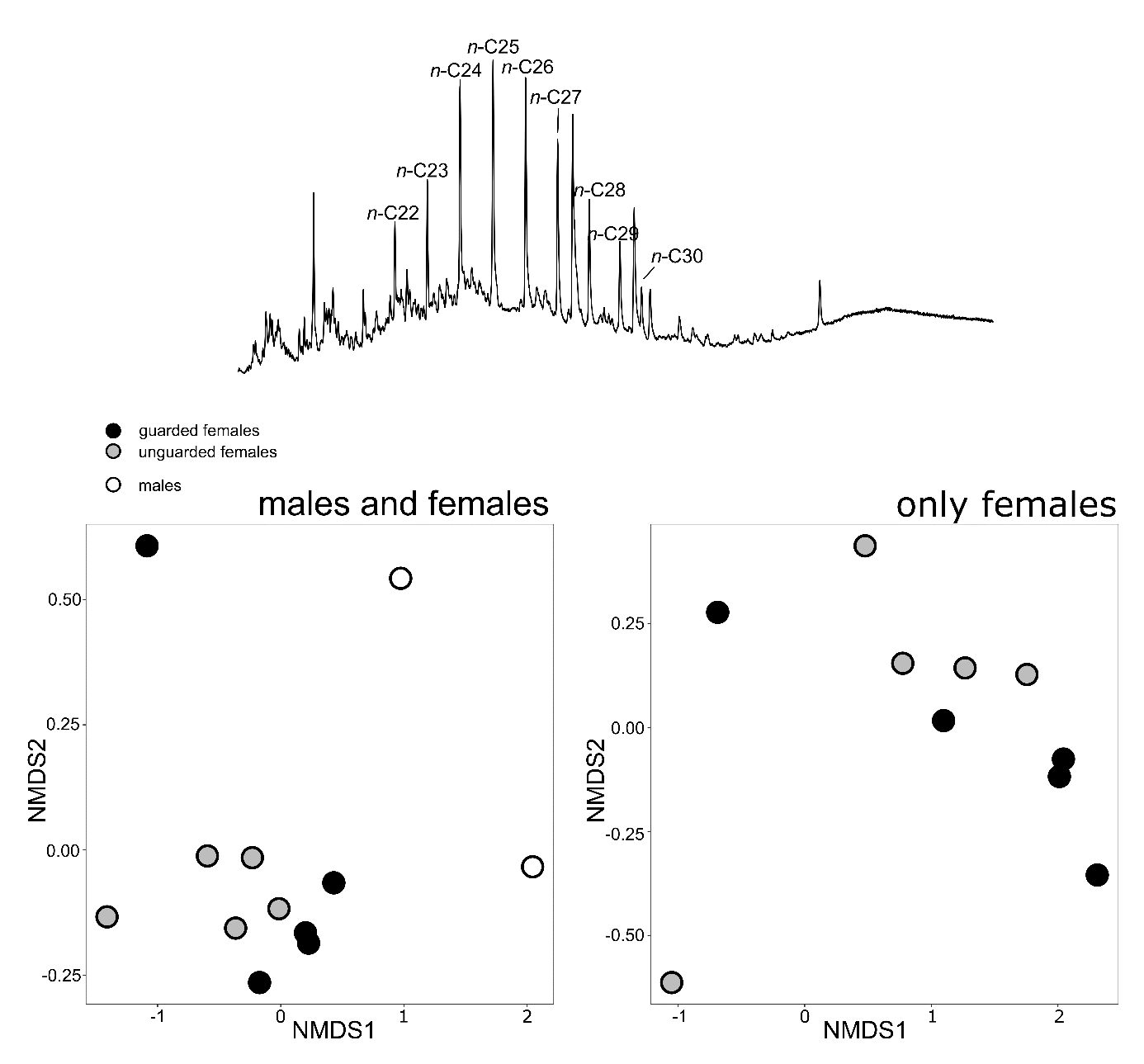
**4**

**Supplementary figure 4:** CHC profiles of representative female *T. urticae*, upper panel. Non-metric multidimensional scaling (NMDS) plot representing showing differences between the males (white circles) but no clear difference between the guarded females (black circles) and unguarded females (grey circles), lower left. NMDS plot showing only the guarded females (black) and unguarded females (gray).
